# A direct selection strategy for isolating aptamers with pH-sensitive binding activity

**DOI:** 10.1101/460063

**Authors:** Chelsea K. L. Gordon, Michael Eisenstein, Hyongsok Tom Soh

## Abstract

An aptamer reagent that can switch its binding affinity in a pH-responsive manner would be highly valuable for many biomedical applications including imaging and drug delivery. Unfortunately, the discovery of such aptamers is difficult and only a few have been reported to date. Here we report the first experimental strategy for generating pH-responsive aptamers through direct selection. As an exemplar, we report streptavidin-binding aptamers that retain nanomolar affinity at pH 7.4 but exhibit a ~100-fold decrease in affinity at pH 5.2. These aptamers were generated by incorporating a known streptavidin-binding DNA motif into an aptamer library and performing FACS-based screening at multiple pH conditions. Upon structural analysis, we found that one aptamer’s affinity-switching behavior is driven by a non-canonical G-A base-pair that controls its folding in a highly pH-dependent manner. We believe our strategy could be readily extended to other aptamer-target systems because it does not require *a priori* structural knowledge of the aptamer or the target.

---

Cellular pH is carefully regulated, as it plays an essential role in many critical functions including energy generation and maintenance of protein structure and function.^1–3^ Additionally, differences in pH help to control the binding and release of important biomolecules by pH-regulated receptors. One critical example is hemoglobin, which exhibits reduced affinity for oxygen as pH decreases.^4^ This promotes uptake of oxygen in the lungs, where the pH is higher, and the subsequent release of oxygen into muscle tissue, where the pH is lower. Reagents that exploit these pH differences for controlled activation or release have proven valuable for many biotechnology applications, most notably in the areas of drug delivery and imaging.^5–8^ For example, several groups have described pH-sensitive DNA nanostructures^9^ that can perform a wide array of molecular functions, such as sequestering a drug in an inactive state until reaching a cellular compartment with a permissive pH environment, or intracellular imaging to measure pH gradients within cells.^7,10–14^

Aptamers are a widely-used class of affinity reagents and several studies have demonstrated the feasibility of introducing a diverse range of specialized functionalities into aptamers.^15,16^ In the context of molecular detection or controlled drug release, it would be especially advantageous to have aptamers for which the affinity is modulated by environmental pH, but only a small number of pH-sensitive aptamers have been reported to date.^17,18^ These were produced by engineering known pH-responsive motifs into existing aptamers. For example, the Ricci group designed a cocaine-binding aptamer that incorporates a pH-dependent triplex, and were able to modulate the affinity of the aptamer for cocaine through pH changes.^17^ The DeRosa group added a polyadenine tail to a thrombin-binding aptamer and found that GA mismatches formed at acidic pH, disrupting the aptamer’s G-quadruplex structure and releasing bound thrombin.^18^ This design method has yielded some useful pH-sensitive aptamers, but is somewhat limited because it requires *a priori* knowledge regarding the structure of active binding motifs within existing aptamers in order to guide the incorporation of pH-responsive elements. Furthermore, this approach is constrained by access to a limited range of known pH-sensitive motifs, which may not necessarily perform optimally—or may even impede binding function—after incorporation into a given aptamer sequence.

In an effort to overcome these limitations, we have devised a strategy that enables the direct selection of aptamers that exhibit both excellent target binding and sensitive pH response. To achieve this, we adapted the particle display platform previously developed by our group, in which nucleic acid libraries are converted into monoclonal aptamer particles^19,20^ that can be rapidly and quantitatively screened using a fluorescence-activated cell sorter (FACS). We have designed a selection procedure that enables us to isolate aptamers that exhibit different target affinities at pH 7.4 compared to pH 5.2, which we demonstrate by generating pH-sensitive aptamers for streptavidin. After only three rounds of screening, we generated an aptamer whose affinity for streptavidin differed by approximately two orders of magnitude between pH 5.2 and 7.4. We also performed structural and mechanistic analysis of one of our aptamers and found that its pH sensitivity is governed in part by a single G-A mismatch, which is known to be stabilized at acidic pH. We believe our strategy could be generalized for generating high-quality pH-sensitive aptamers for a wide range of other molecules, eliminating much of the labor and constraints associated with conventional aptamer design strategies. Such aptamers would useful for both *in vivo* and *in vitro* applications, including drug delivery, sensing, and the development of pH-sensitive smart nanomaterials.

## Results and Discussion

### Library design and screening strategy

We devised a molecular library design that enabled us to screen directly for aptamers that exhibit pH-dependent target binding. Each library molecule comprises a known aptamer sequence fused to a 20-nucleotide (nt) randomized domain (**Fig. 1A**), with these two segments flanked by PCR primer-binding sites. The objective is to isolate library molecules that maintain a conformation favoring target binding at permissive pH, but which undergo denaturation or refolding at non-permissive pH into a conformation that subsequently promotes target release. We chose streptavidin as a model target because it is a well-characterized protein that remains stable across a wide pH range. We chose to use SBA29, which was previously isolated by Bing and coworkers and has a reported *K*_*d*_ of 40 ± 18 nM.^21^

**Figure 1.**
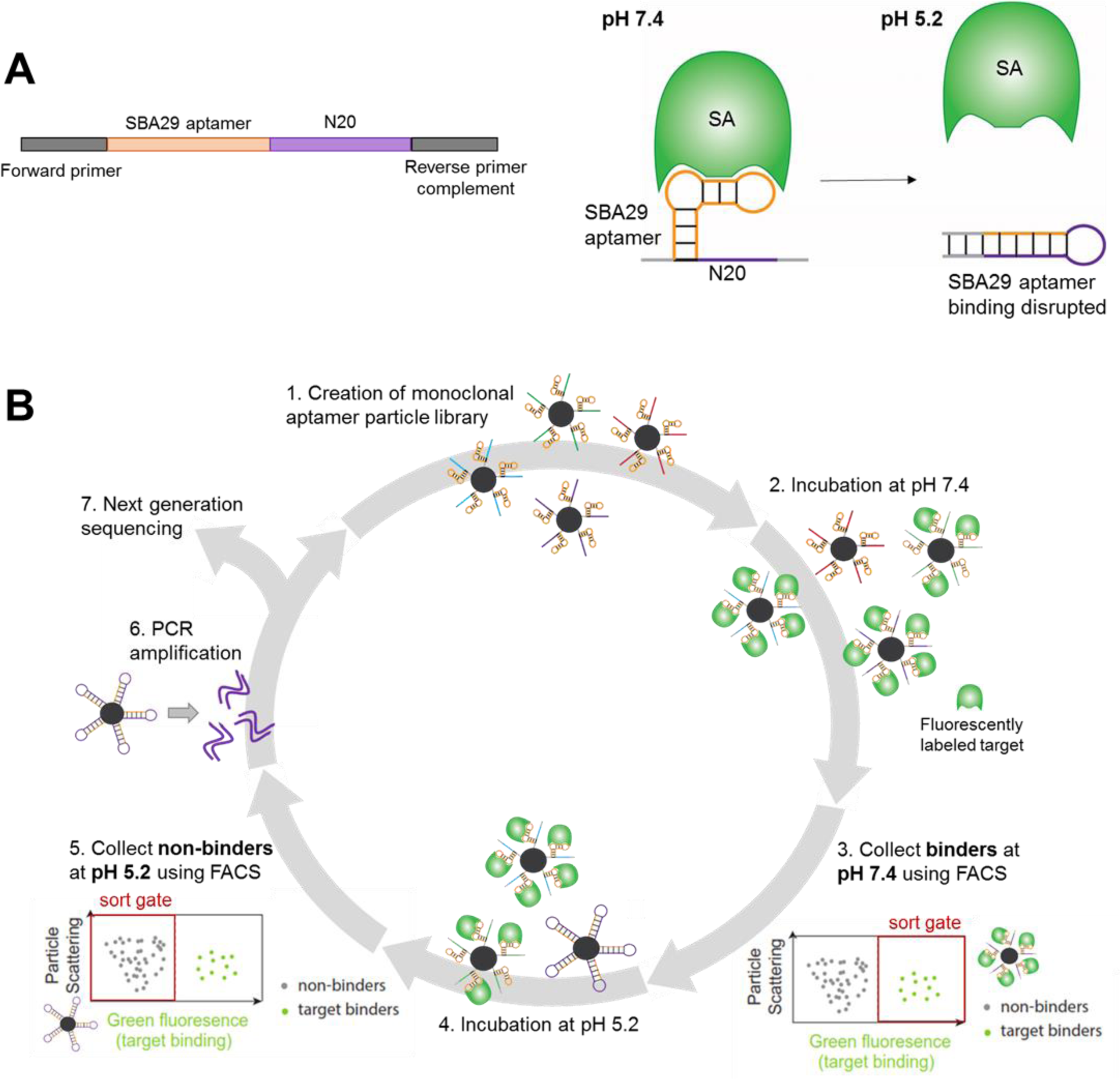
Overview of pH-based particle display screening. (A) Our library design includes a known aptamer sequence—in this demonstration, the streptavidin aptamer SBA29—and a 20-nt random region. The objective of this screen was to identify sequences that bind streptavidin at pH 7.4 but experience disruption of the SBA29 aptamer domain at pH 5.2 to eliminate target binding. (B) Scheme for pH-switching particle display screen. 1) Emulsion PCR is used to generate monoclonal aptamer particles, which are then 2) incubated with fluorescently-labeled streptavidin at pH 7.4. 3) FACS is used to collect aptamer particles that are bound to the fluorescent target at this pH. 4) These are then incubated with labeled streptavidin at pH 5.2, and 5) FACS is used to collect only non-fluorescent aptamers, which no longer bind the target at this acidic pH. These are either 6) subjected to PCR amplification for another round of screening or 7) sequenced for further characterization.

We screened for aptamers that maintain target binding at pH 7.4 but release their target at pH 5.2. Our procedure (**Fig. 1B**) is a variation on the previously-described particle display platform, a high-throughput aptamer screening strategy based on FACS that enables the analysis of individual aptamer binding characteristics at a rate of ~10^6^ sequences/hour.^19^ The critical difference between this platform and conventional SELEX (systematic evolution of ligands by exponential enrichment) is that aptamers in solution are transformed into monoclonal aptamer particles, allowing the measurement and sorting of each individual aptamer sequence. FACS enables fine discrimination of the highest affinity aptamers in a given pool, leading to much higher enrichment rates in each round of screening than are possible with conventional SELEX.

First, we used emulsion PCR to convert a solution-phase DNA library of ~10^9^ molecules into monoclonal aptamer particles, which each display many copies of a single sequence (**Fig. 1B**, step 1). These were then subjected to two sequential sorting procedures to identify sequences with pH-dependent binding behavior. For the first sort, we incubated the aptamer particles with fluorescently-labeled streptavidin in pH 7.4 selection buffer (step 2) and used FACS to collect all particles that bound streptavidin at this pH (step 3). We then sought to isolate aptamer particles that lost their ability to bind streptavidin under more acidic conditions, and so we re-incubated the collected particles with streptavidin in pH 5.2 selection buffer (step 4). In the subsequent round of FACS, we collected all non-fluorescent aptamer particles, representing sequences that could no longer bind streptavidin as a result of a pH-induced conformational change (step 5). The aptamer particles collected at this step were PCR amplified to create the aptamer pool for the next round (step 6). After three rounds of screening, we subjected all three aptamer pools to high-throughput sequencing (Step 7).

### Particle display screening for pH-switching aptamers

We performed three rounds of particle display screening against streptavidin, as described above. Prior to each sorting step, we incubated the aptamer particles with 200 nM streptavidin labeled with Alexa Fluor 488 (SA-AF488) for 1 hour. In the first FACS screen, all of the sequences that bound streptavidin at pH 7.4 were collected. We defined the non-fluorescent reference gate using unlabeled beads, and defined the sort gate to include all sequences with higher fluorescence intensity than this background level. We did not set the gates to exclude lower affinity aptamers because we wanted to optimize our selection for the identification of aptamers that undergo pH-induced switching rather than selecting primarily for extremely high affinity to streptavidin. We then incubated the collected aptamer particles from the first FACS sort step with 200 nM SA-AF488 in pH 5.2 selection buffer for 30 minutes. For the second sorting step, we collected all of the aptamer particles that did not bind streptavidin at pH 5.2. Sorting was performed with the same gate positions as in the first sorting step, but this time we collected the aptamer particles from the reference (non-binding) gate and discarded those in the high fluorescence gate. The second round was performed with the same conditions as the first round. In the third round, only the first FACS sort step was performed, collecting aptamer particles with high fluorescence intensity after incubation with 200 nM SAAF488 at pH 7.4.

Over the course of three rounds, we observed a clear increase in both streptavidin binding and pH-induced switching behavior of the aptamer pool (**Fig. 2**). After preparing the aptamer particles for each pool, we tested the binding of the particles to streptavidin at pH 7.4 and pH 5.2 as a prelude to particle display screening. Based on this analysis, we determined that the proportion of aptamer particles residing within the sort gate increased from 3.1% of the initial library to 11.8% of the round 3 pool, indicating a clear increase in the number of streptavidin-binding sequences. More notably, the proportion of aptamer particles that retained binding to their target at pH 5.2 steadily decreased over the course of screening, from 6.92% in round 1 to 5.05% in round 2 to just 1.35% in the final round. Based on these measurements, we determined that the ratio of binding at pH 7.4 to binding at pH 5.2 increased from 1.1 for the starting library to 2.1 and 1.9 for the round 1 and round 2 pools, respectively. For the round 3 pool, this ratio increased dramatically to 8.7. As the round 3 pool demonstrated strong pH-sensitivity, we did not perform further rounds of screening.

**Figure 2.**
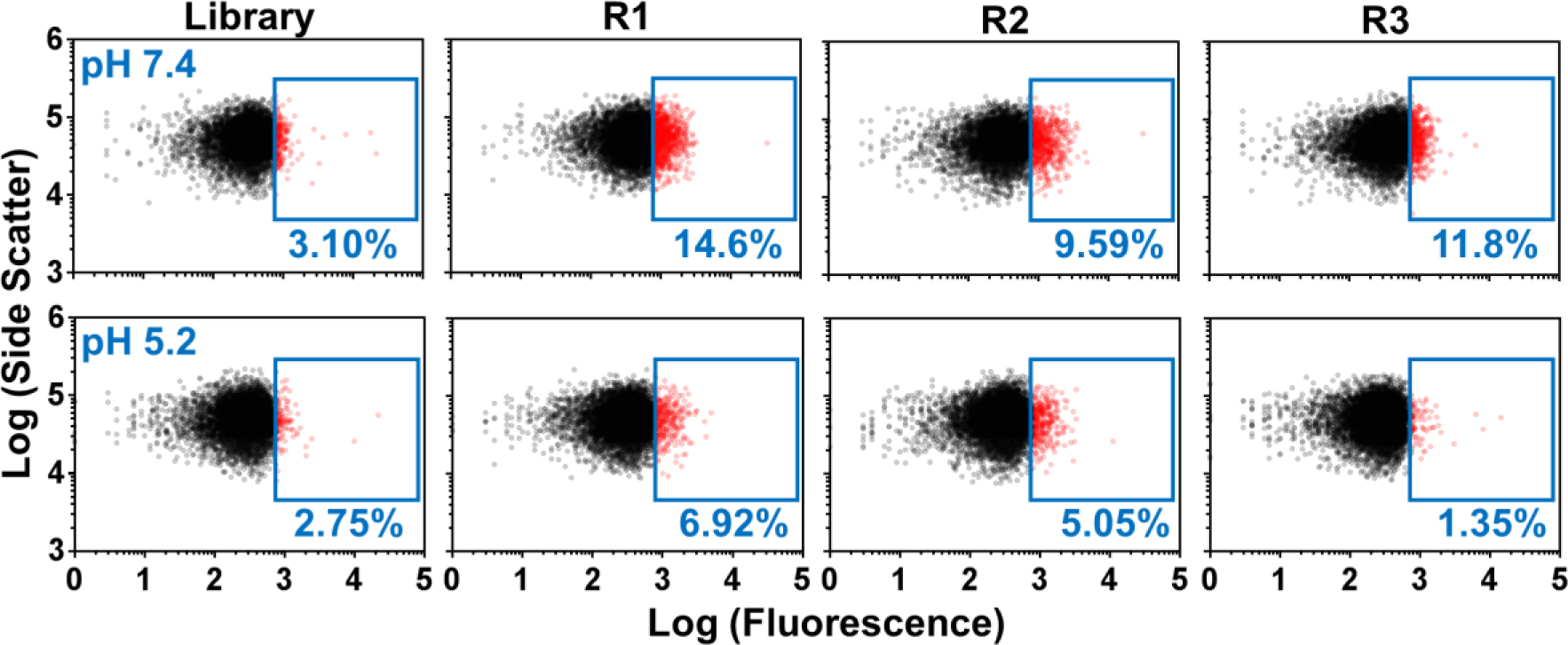
Binding assays for pools from each round of our pH-dependent particle display screen. Each set of binding measurements was collected prior to screening. The box denotes aptamer particles with fluorescence above background at pH 7.4 (top) and pH 5.2 (bottom). The percentage of particles residing within the high fluorescence gate is shown in each plot. By round 3, a large population of aptamer particles exhibited pH-switching, with high binding at pH 7.4 and low binding at pH 5.2.

### High-throughput sequencing reveals aptamers enriched based on pH sensitivity

To better characterize the enrichment that had taken place, we performed high-throughput sequencing of the three aptamer pools. We prepared the pools for sequencing by adding different indices to each pool using the Nextera XT DNA Library Preparation Kit from Illumina (see **SI**). Sequencing was performed using an Illumina MiSeq at the Stanford Functional Genomics Facility. After filtering out low-quality sequences, we obtained 1,035,183 reads with 363,155 unique sequences (35.1%) in round 1, 1,160,070 reads with 257,612 unique sequences (22.2%) in round 2, and 1,150,478 reads with 180,777 unique sequences (15.7%) in round 3. This indicates that even though the diversity of the pool decreased each round, there was still considerable diversity in the round 3 pool. We analyzed the copy number and enrichment of each sequence to identify top aptamer candidates for further functional characterization, and identified several aptamers that greatly outperformed the rest of the pool in terms of either their copy number in round 3 or in their enrichment from round 1 to round 3 (**Fig. 3A**). We selected ten sequences for further analysis: the seven most highly-enriched sequences and three most abundant sequences from round 3 (**Table S-1**).

We synthesized these ten candidate sequences and examined their binding characteristics in a fluorescence assay, incubating particles displaying each sequence with a streptavidin-phycoerythrin (SA-PE) conjugate (**Fig. 3B**). Eight of the ten sequences exhibited greater binding to streptavidin at pH 7.4 than at pH 5.2, and we selected the two sequences that showed the largest decrease in binding from pH 7.4 to pH 5.2 (S3 and S8) for further testing.

**Figure 3.**
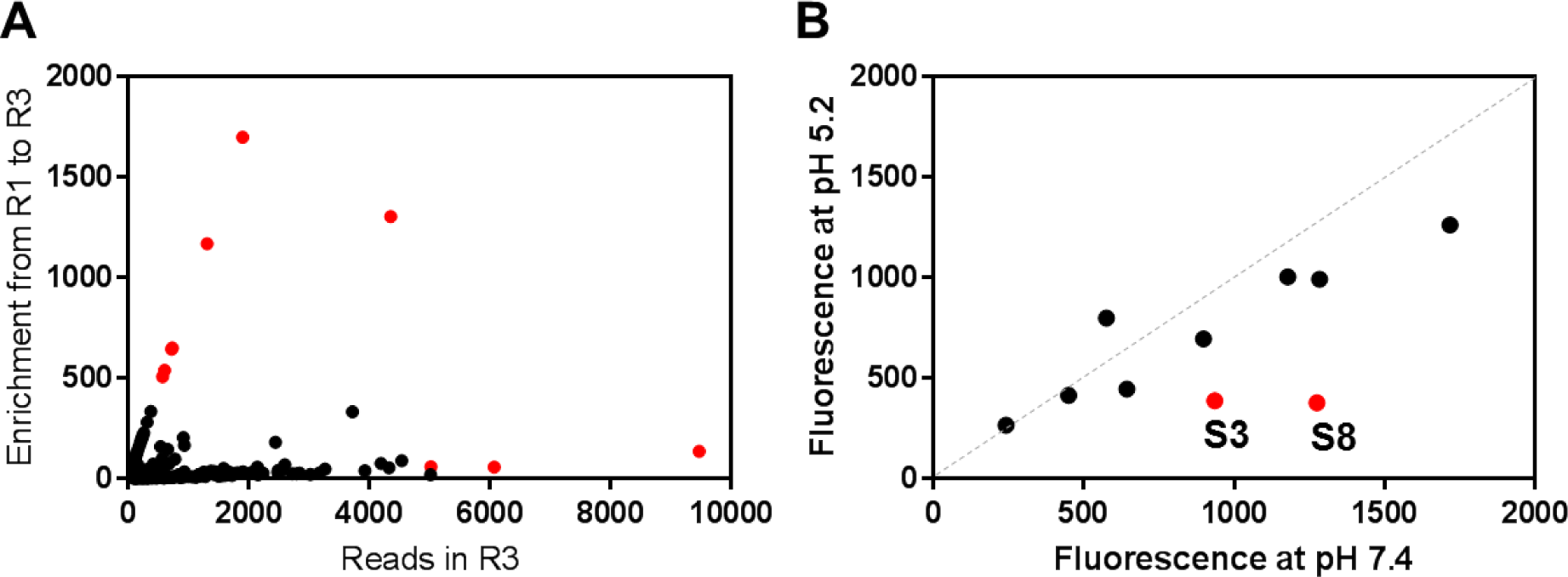
Identification of pH-responsive aptamer candidates. (A) Plot shows the 1,000 most abundant sequences from round 3 of screening after filtering out low-quality reads and sequences with incorrect length. Red dots show the seven most highly-enriched sequences from round 1 to round 3 (upper left) and the three most abundant sequences from round 3 (lower right), which were selected for further testing. (B) These sequences were tested for pH-dependent binding in a fluorescence assay. Each sequence was conjugated to beads, and binding to streptavidin-phycoerythrin (SA-PE) conjugates (50 nM) was measured at pH 7.4 and pH 5.2. We selected the two sequences with the greatest difference in SA-PE binding at pH 7.4 and pH 5.2 (shown in red) for further characterization.

### Aptamers isolated via particle display exhibit strong pH sensitivity

Our analysis of S3 and S8 revealed that our selection procedure is highly effective at isolating pH-responsive derivatives of existing aptamers. We generated particles displaying these two sequences as well as the original SBA29 aptamer and used flow cytometry to measure the fluorescence intensity of the aptamer particles after incubating with SA-PE at a range of concentrations at both pH 7.4 and pH 5.2 (**Fig. 4**). We used a saturation binding model (one-site, total binding) to determine the equilibrium dissociation constant (*K*_*d*_) for each sequence. SBA29 exhibited minimal pH sensitivity, with a similar *K*_*d*_ under both conditions: 10.4 ± 1.5 nM at pH 7.4 and 3.50 ± 0.46 at pH 5.2.

**Figure 4.**
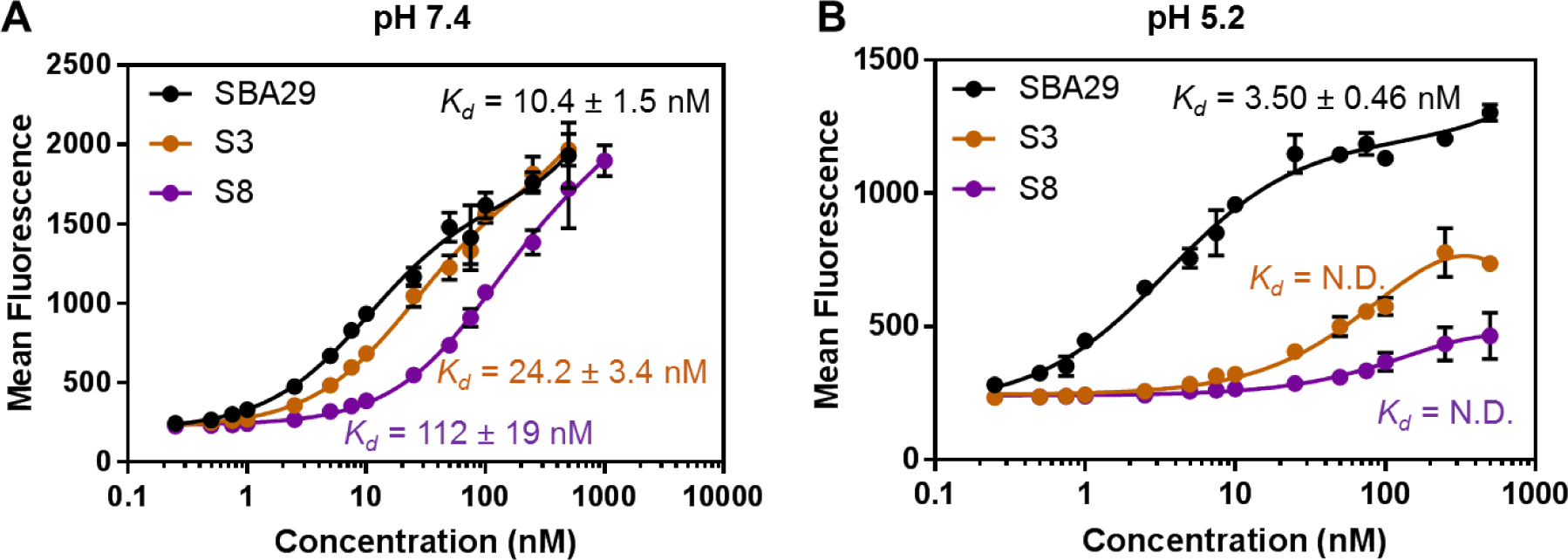
Streptavidin-binding measurements of SBA29 and the selected aptamers S3 and S8 in a fluorescent bead-based assay at (A) pH 7.4 and (B) pH 5.2. Error bars were determined from the standard deviation of experimental replicates (n = 2 for SBA29, pH 5.2, n = 3 for all other samples). (N.D. = not determined)

In contrast, the binding affinity was strongly pH-dependent for both S3 and S8. These two aptamers bound strongly to streptavidin at pH 7.4, with a *K*_*d*_ of 24.2 ± 3.4 nM and 112 ± 19 nM for S3 and S8, respectively. However, both aptamers had much weaker binding at pH 5.2. Indeed, we were not able test high enough target concentrations to reach a stable bound plateau for either aptamer at pH 5.2 in order to obtain a meaningful *K*_*d*_ (**Fig 4B**). As a control experiment, we also measured the fluorescence of forward primer-conjugated beads without aptamers at both pH values, with and without SA-PE. We observed minimal signal, demonstrating that non-specific target-bead interactions do not produce any meaningful background at either pH 5.2 or 7.4 (**Fig. S-1**).

We chose to perform more detailed characterization for S8 because it had minimal binding at pH 5.2, indicating strong pH-sensitivity. Since bead-based fluorescent measurements are performed with many aptamers conjugated to particles, avidity effects can impact the measured binding affinity. We therefore used microscale thermophoresis (MST) to independently assess the solution-phase binding affinity of SBA29 and S8. As with our bead-based assay, MST demonstrated the pH-insensitivity of SBA29, which exhibited a *K*_*d*_ of 6.1 nM and 27 nM at pH 7.4 and at pH 5.2, respectively (**Fig. 5A**, **B**), whereas S8 again exhibited striking pH sensitivity. At pH 7.4, we determined that S8 has a *K*_*d*_ of 10 nM (**Fig. 5C**); this is ~10-fold lower than the *K*_*d*_ we measured by bead-based measurements, but represents reasonable agreement given the differences in the two measurement techniques. But at pH 5.2, as with the bead-based assay, S8’s affinity was too low to obtain a meaningful *K*_*d*_ (**Fig. 5D**). From the observed binding response, we estimate that the *K*_*d*_ is in the high nanomolar to low micromolar range, which indicates that our aptamer’s streptavidin affinity at pH 7.4 is roughly two orders of magnitude higher than at pH 5.2. In order to better characterize the nature of S8’s pH response, we measured streptavidin binding at a range of pH values between pH 5.2 and pH 7.4. This yielded a sigmoidal binding curve, indicating a gradual rather than single-step pH response, where half maximal signal occurs at pH 6.5 (**Fig. S-2**).

**Figure 5.**
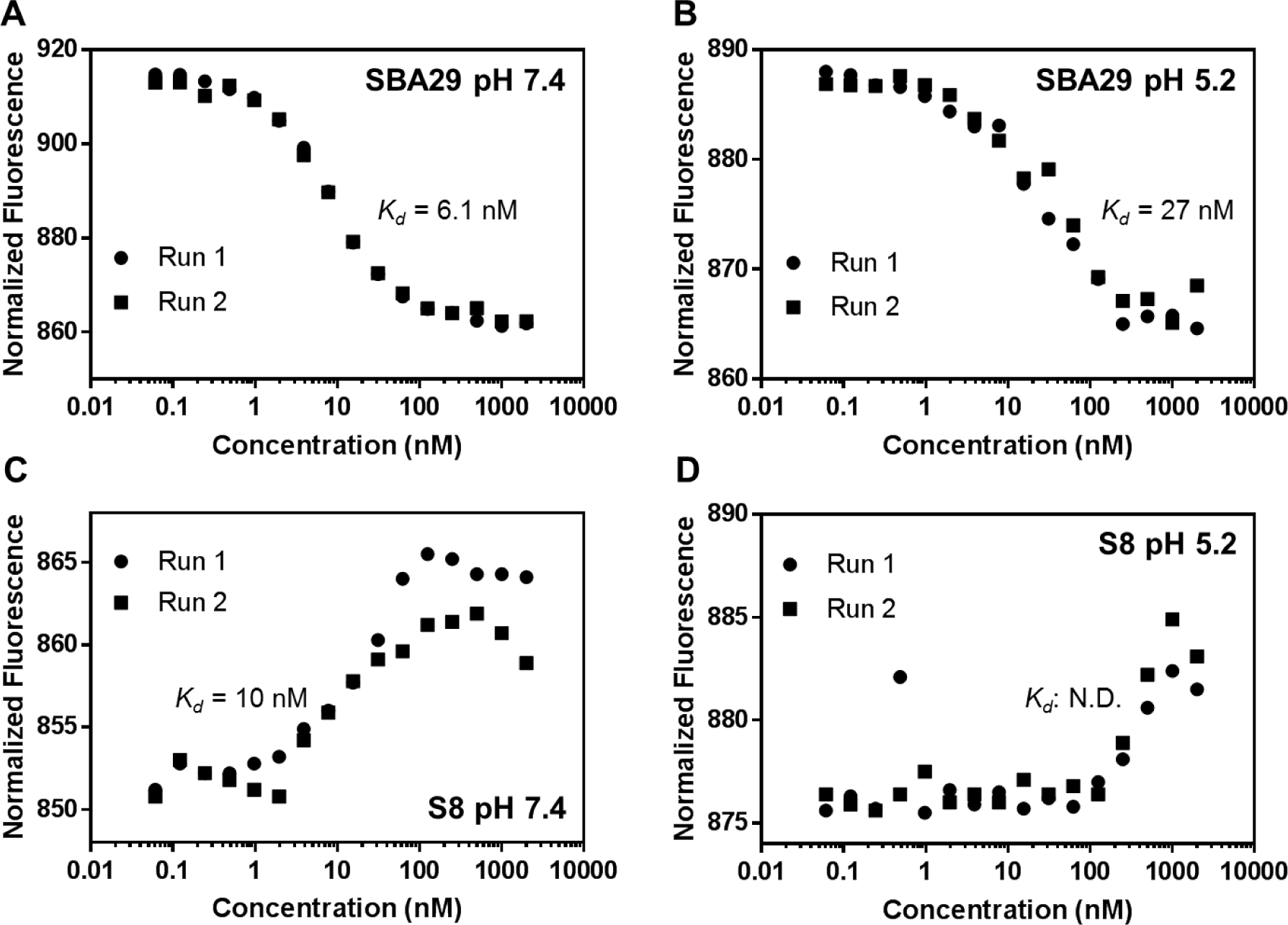
Binding measurements by microscale thermophoresis for SBA29 at (A) pH 7.4 and (B) pH 5.2 and for S8 at (C) pH 7.4 and (D) pH 5.2. *K*_*d*_ is shown for all experiments except S8 at pH 5.2, for which this measurement could not be determined reliably.

### A nucleotide mismatch contributes to pH sensitivity

After affinity testing, we predicted the secondary structure for S8 using mfold.^22^ Our analysis determined that S8 has two predicted secondary structures. In one, SBA29 retains its nominal conformation, with the randomized domain hybridized to one of the primer-binding sequences (**Fig. 6A**). In the second structure, the randomized region hybridizes with the SBA29 sequence, preventing it from folding into a conformation that enables streptavidin binding (**Fig. 6B**). The base-pairing between SBA29 and the randomized domain within the latter, ‘blocked’ structure contains a predicted G-A mismatch, a pairing which has been computationally and experimentally shown to be stabilized at acidic pH.^23–25^ We therefore hypothesized that the latter structure may be energetically favorable at pH 5.2, whereas the first structure, in which SBA29 is properly folded, is more stable at pH 7.4.

**Figure 6.**
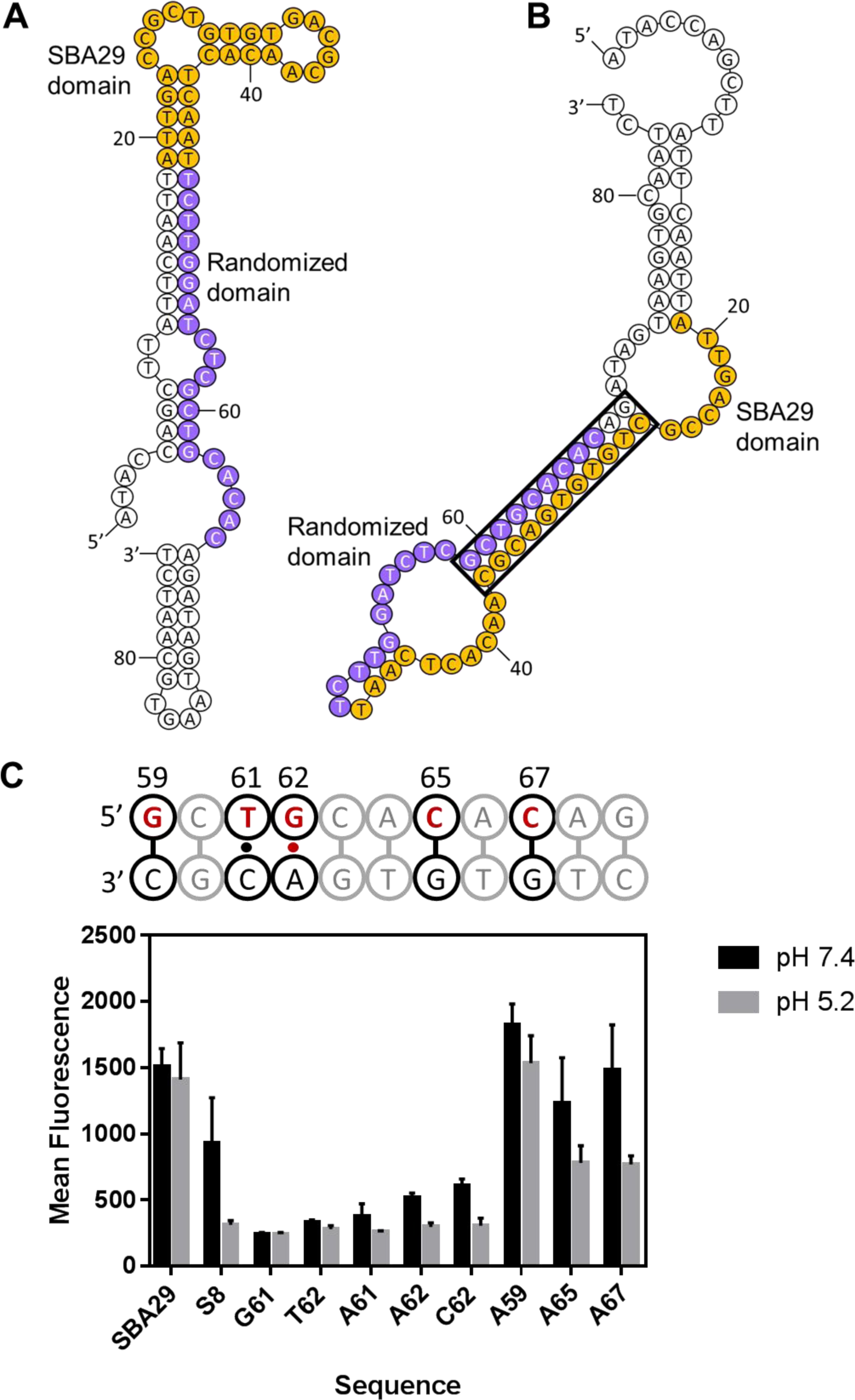
Predicted secondary structures for pH switching aptamer S8. (A) The SBA29 aptamer domain (orange) is folded correctly in the lowest free energy structure and does not interact with the randomized domain (purple). (B) Another predicted low free energy structure for the same sequence (right) shows the SBA29 domain (orange) blocked by the randomized domain (purple). The portion of the sequence shown in (C) is outlined in black. (C) (top) Positions of mutated bases are shown in red. The pH-sensitive G-A mismatch predicted to stabilize the blocked structure is shown as a red dot. The C-T mismatch is shown as a black dot. Watson-Crick base pairs are shown as dashes. (C) (bottom) Bead-based binding assay of S8 mutant sequences at 50 nM streptavidin. Three experimental replicates were performed, and mean + SD is shown.

We generated various point-mutations in the S8 aptamer that were predicted to affect the stability of the low-pH blocked structure (red bases in **Fig. 6C)**. We then tested these S8 variants in a binding assay in which we incubated aptamer particles displaying each mutant sequence with fluorescently labeled streptavidin at pH 7.4 and pH 5.2. By measuring the fluorescence intensity of the streptavidin-bound aptamer particles, we were able to identify which mutations increased or decreased the affinity of the aptamer to streptavidin at each pH. First, we replaced the mismatches at positions 61 and 62 with nucleotides that enable canonical base pairing (G61 and T62); unlike the G-A pairing at base 62, the predicted C-T mismatch that normally occurs at base 61 is not stabilized at acidic pH.^23^ We expected that these substitutions would stabilize the blocked structure, and indeed, these two mutations both exhibited greatly reduced binding (by 75% and 65%, respectively) at pH 7.4.

Next, we introduced other mismatches at the predicted pH-dependent mismatch site. Based on mfold simulations, both the A62 and C62 variants are predicted to favor the blocked structure (ΔG = −23.52 kcal/mol and −23.22 kcal/mol, respectively). Although these structures are slightly less stable than G61 and T62 (ΔG = −27.77 kcal/mol and −25.08 kcal/mol, respectively), both sequences still have significantly reduced streptavidin binding at pH 7.4. Notably, replacing the pH-sensitive G-A mismatch with a C-A mismatch (C62) significantly reduced binding at pH 7.4, even though C-A is also selectively stabilized at acidic pH.^23,25^ This shows that the G-A mismatch provides the correct balance to favor folding of SBA29 at pH 7.4 and to disrupt this binding by stabilizing the blocked structure at pH 5.2.

Finally, we replaced three different G-C pairs in the stem of the blocked structure with a C-A or G-A mismatch (A59, A65, A67) to see if the introduction of a second pH-sensitive mismatch would strengthen S8’s pH-switching behavior. We observed far less pH responsiveness in A59, with high levels of binding in both pH conditions. We hypothesize that this is because the elimination of the G-C pair greatly destabilizes the stem of the blocked structure and enables the SBA29 domain to remain folded at both pH values. The introduction of a second G-A mismatch to the stem of the blocked structure in A65 and A67 enabled retention of high binding at pH 7.4, but also resulted in moderately high levels of binding at pH 5.2. This is likely because the G-C pair is more stable than the G-A mismatch, even at acidic pH, such that the stem in the blocked structure becomes less stable. Nevertheless, these sequences still retained some pH sensitivity.

Overall, these results support a model in which hybridization between the randomized domain in S8 and the SBA29 aptamer domain contribute to the formation of a ‘blocked’ structure that is incapable of binding to streptavidin. Although the full mechanism of this pH-switching behavior is presently not fully understood, the pH-dependence of the non-canonical G-A pairing at site 62 appears to play a critical role in determining aptamer stability and conformation at acidic versus neutral pH conditions.

## Conclusion

In this work, we describe a rapid and high-throughput method that enables us to screen for pH-sensitive derivatives of existing aptamers based on particle display, without the need for labor-intensive aptamer engineering procedures.^10,26^ As a demonstration, we isolated aptamers that exhibit high affinity for streptavidin at neutral pH but release their cargo under acidic conditions after only three rounds of screening. One of these aptamers, S8, retained the nanomolar target affinity of its parent aptamer at pH 7.4, but exhibited an estimated 100-fold decrease in streptavidin affinity at pH 5.2 versus pH 7.4. Upon modeling the predicted secondary structure of S8, we identified two different conformations for this aptamer that appear to be governed in part by a pH-sensitive, non-canonical base-pair. At neutral pH, the streptavidin-binding aptamer domain retains the secondary structure of the non-pH-responsive parent aptamer, SBA29. However, acidic conditions favor a reorganization of the aptamer in which this target-binding domain is incorporated into a stem-loop by base-pairing with the randomized sequence that was selected during our screening process. This stem contains a G-A mismatch with known pH-responsive characteristics, and we used mutational analysis to confirm that both this base-pair and the stem-forming elements of the randomized domain in general are critical to the aptamer’s pH-responsive characteristics. These results demonstrate that our screening method can be used to generate high-affinity aptamers with pH-responsive functionality without relying exclusively on previously-identified pH-sensitive motifs.^17,18^ As such, we believe this approach will prove highly valuable for generating environmentally-responsive aptamers for drug delivery, biosensors, and a variety of other applications.

---

For TOC only

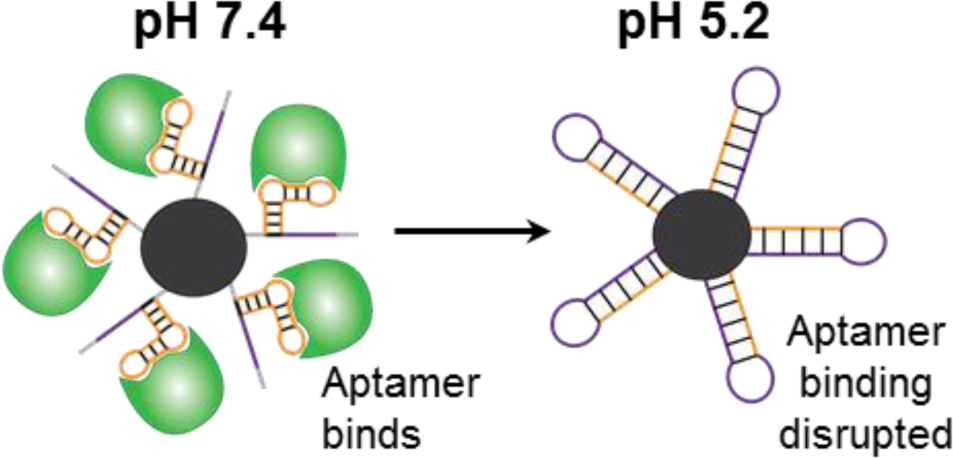

